# Peptide collision cross sections of 22 post-translational modifications

**DOI:** 10.1101/2022.12.23.521814

**Authors:** Andreas Will, Denys Oliinyk, Florian Meier

**Affiliations:** Functional Proteomics, Jena University Hospital, 07747 Jena, Germany

**Keywords:** post-translational modifications, PTM, ion mobility, TIMS, collision cross section, CCS

## Abstract

Recent advances have rekindled the interest in ion mobility spectrometry as an additional dimension of separation in mass spectrometry (MS)-based proteomics. It separates ions according to their size and shape in the gas phase. Here, we set out to investigate the effect of 22 different post-translational modifications (PTMs) on the collision cross section (CCS) of peptides. In total, we analyzed ∼4700 pairs of matching modified and unmodified peptide ions by trapped ion mobility spectrometry (TIMS). Linear alignment based on spike-in reference peptides resulted in highly reproducible CCS values with a median coefficient of variation of 0.3%. On a global level, we observed a redistribution in the *m/z* vs. ion mobility space for modified peptides upon changes in their charge state. Pairwise comparison between modified and unmodified peptides of the same charge state revealed median shifts in CCS between – 1.1% (lysine formylation) and +4.5% (O-GlcNAcylation). In general, increasing modified peptide masses were correlated with higher CCS values, in particular within homologous PTM series. However, investigating the ion populations in more detail, we found that the change in CCS can vary substantially for a given PTM depending on the gas phase structure of its unmodified counterpart. In conclusion, our study shows PTM- and sequence-specific effects on the cross section of peptides, which could be further leveraged for proteome-wide PTM analysis.

## Introduction

Post-translational modifications (PTMs) are key regulators of protein activity and function in health and disease. Not at least because most PTMs involve a specific shift in the molecular weight of modified amino acids, mass spectrometry (MS)-based proteomics has evolved as the method of choice for the investigation of PTMs on a proteome-wide scale (1-3). MS-based proteomics analyzes complex mixtures of (modified) peptides derived from tryptic digests by liquid chromatography coupled to high-resolution MS. Since the modification mass shift also transfers to fragment ions, this allows identifying modified peptide sequences and also often localizing the PTM to a specific amino acid.

Recently, ion mobility spectrometry (IMS) has become a popular extension of the proteomics toolbox that adds one more dimension of separation (4-7). IMS distinguishes ions in the gas phase by their size and shape, which can be inferred from ion mobility measurements in the form of a collision cross section (CCS) or, more precisely, the momentum transfer collision integral. (8-10). In the CCS vs *m/z* dimension, electro sprayed tryptic peptides typically split into distinct populations according to their charge state (11). A more fine-structured heterogeneity, most prominently for triply charged species, has been attributed to either more extended or more compact structures that are determined by the linear peptide sequence (12, 13). As a result, even isobaric and isomeric peptides can have distinct cross sections (14, 15).

Trapped ion mobility spectrometry (TIMS) is a relatively new type of IMS that inverses the concept of classical drift tube IMS by holding ions in an electric field against an opposing gas flow (16-18). Lowering the electric field strength releases ions sequentially from the TIMS device to the downstream mass analyzer as a function of increasing ion mobility (or decreasing CCS). The PASEF acquisition mode synchronizes the separation with a quadrupole time-of-flight mass analyzer, which greatly enhances the speed and sensitivity of peptide sequencing (5, 19, 20). We have recently shown that an intriguing feature of this setup is that it enables peptide CCS measurements at very large scale and with high precision, sufficient to train a deep learning model to predict peptide cross sections based solely given the linear amino acid sequence and charge state as an input (13).

The advantages of IMS in terms of speed, sensitivity and specificity should equally apply to or even be enhanced in the analysis of PTMs, given the additional complexity arising from isobaric modifications and positional isomers (21-24). This motivated researchers since the beginnings of IMS to study the effect of specific modifications on the gas phase characteristics of model peptides. By far the most studied example is phosphorylation of serine and threonine residues, which, interestingly, often leads to more compact gas phase structures compared to unmodified peptides – despite the increase in mass (23, 25-27). Another field of interest are N-glycosylated peptides, for which the bulky glycan moiety results in a distinct separation from unmodified peptides in the ion mobility dimension, and even differentiation between isomeric localization variants has been demonstrated. (24, 28, 29). To model the effect of PTMs on peptide CCS values, Clemmer and co-workers generated a data set of cysteine-palmitoylated and -carboxyamidomethylated peptides and derived modification-specific intrinsic size parameters (30). Kaszycki and Shvartsburg extended this approach to predict the intrinsic size parameters of 100 different PTMs (31). More recent large-scale studies further highlighted additional sequence-dependent effects on peptide cross sections resulting from intra-molecular interactions (13, 23, 32).

The further exploration of the potential of IMS for PTM analysis and the development of more accurate prediction models for a wide range of PTMs, similar to those for unmodified peptides (13, 32, 33), is currently limited by the availability of comprehensive experimental data. Here, we set out to investigate the effect of 22 different naturally-occurring PTMs on the collision cross section of peptides, including 21 sets of modified peptides synthesized as part of the ProteomeTools project (34, 35) and an additional set of O-GlcNAcylated peptides.

## Methods

### Synthetic peptides

A library of ∼5,000 lyophilized synthetic peptides in 96-well format representing 21 naturally occurring post-translational modifications on K, R, P and Y residues and their unmodified counterparts were obtained from the ProteomeTools project (35). Additionally, we purchased synthetic O-GlcNAcylated peptides and corresponding unmodified peptides from JPT Peptide Technologies GmbH (SpikeMix PTM-Kit 57 and 55). The pre-pooled peptides were reconstituted in 2% acetonitrile/0.1% formic acid to a final concentration of ∼100 fmol/µL. As a reference, we spiked each sample with a retention time standard (Biognosys iRT) in a ratio of 1:40 (vol/vol).

### Liquid chromatography and mass spectrometry

Nanoflow reversed-phase liquid chromatography was performed on a nanoElute system (Bruker Daltonics). Peptides were separated with a 120 min gradient at a flow rate of 0.3 µL/min at 60 °C on a homemade 50 cm x 75 µm column with a pulled emitter tip, packed with 1.9 μm ReproSil-Pur C18 - AQ beads. Mobile phases A and B were water/0.1% formic acid and acetonitrile/0.1% formic acid. The LC system was connected online to a TIMS quadrupole time-of-flight mass spectrometer (Bruker timsTOF Pro) via a CaptiveSpray nano-electrospray source (5). TIMS analysis was performed in a range from 1/K_0_ = 1.5 to 0.6 Vs cm^-2^, while accumulating and analyzing incoming ions in parallel for 100 ms. The ion mobility gas was nitrogen from ambient air without temperature control, and the pressure at the TIMS entrance was kept at ∼2.7 mbar. The TIMS elution voltages were externally calibrated to *1/K*_*0*_ values using a linear model fitted to at least three ions from the Agilent ESI LC/MS tuning mix (*m/z, 1/K*_*0*_: 622.0289, 0.9848 Vs cm^-2^; 922.0097, 1.1895 Vs cm^-2^; and 1221.9906, 1.3820 Vs cm^-2^). All data were acquired in dda-PASEF mode and suitable precursor ions for fragmentation were selected by their relative position in the *1/K*_*0*_ vs. *m/z* plane. The quadrupole isolation window was set to 2 Th for *m/z* < 700 and 3 Th for *m/z* > 700. To prevent excessive re-fragmentation, we defined an active exclusion time of 0.4 min.

### Data processing

The mass spectrometry data were processed with MaxQuant (version 2.0.2.0 or 2.1.4.0) using the pre-defined parameters for the analysis of timsTOF data (36, 37). MS/MS spectra were searched against either concatenated peptide sequences as provided by the ProteomeTools consortium or separated peptide sequences as provided by JPT, both supplemented with the iRT peptide sequences. The digestion mode was set to ‘specific’ according to the cleavage rules for Trypsin. We analyzed each PTM pool separately, defining only the respective modification as a variable modification (Supplementary Table 1) and cysteine carbamidomethylation as a fixed modification. The false discovery rate on the peptide spectrum match and protein level was controlled by a target-decoy approach <1%. For modified peptides, we required a minimum Andromeda score (38) of 40 and a minimum delta score of 6 (39).

### Bioinformatic analysis

Data analysis and visualization was performed in R (v4.2.1) using the packages tidyverse (1.3.1), plyr (1.8.7), magrittr (2.0.3), data.table (1.14.2), ggplot2 (3.3.6), ggrepel (0.9.1), ggnewscale (0.4.8), heatmaply (1.4.0), orca (1.1.1), GGally (2.1.2), Peptides (2.4.4) and VennDiagram (1.7.3). Heatmaps were generated with the R package ‘heatmaply’ and clustered using the Euclidean distance measure and the average linkage function.

To account for time-dependent drifts in measured CCS and retention time (RT) values, we aligned all experiments by subtracting a run-specific alignment factor derived from the median deviation of the 11 iRT peptides relative to the median of three reference experiments (Supplementary Table 2). For further analysis, we kept only peptide sequences identified with a single modification site not localized on the C-terminus. In cases, in which MaxQuant listed multiple ‘evidences’ for a modified peptide sequence, we retained only the most abundant feature (highest intensity value) for charge state 2 and 3 for our analysis. CCS and RT values represent median values calculated from one to three technical replicates. Differences in the collision cross section of modified peptides and their unmodified counterparts were calculated relative to the cross section of the unmodified peptide (ΔCCS = (CCS_modified_ - CCS_unmodified_)/ CCS_unmodified_).

### Data availability

The mass spectrometry proteomics data underlying this study have been deposited to the ProteomeXchange Consortium (40, 41) via the partner repository with the dataset identifier PXD038888.

## Results and Discussion

### Constructing a peptide CCS dataset for 22 PTMs

To generate a high-quality CCS dataset of a wide range of naturally occurring peptide modifications, we analyzed pooled libraries of synthetic peptides by nanoflow reversed-phase liquid chromatography and TIMS - quadrupole time-of-flight mass spectrometry (**Fig. 1a**). In dda-PASEF mode, the mass spectrometer selects suitable precursor ions from a survey TIMS-MS scan and targets them for fragmentation in the subsequent PASEF-MS/MS scans. As quadrupole and collision cell are positioned downstream of the TIMS analyzer, precursor and fragment spectra are linked through their position in the ion mobility spectrum. We processed this data in the MaxQuant software to assemble three-dimensional features in ion mobility, *m/z* and retention time dimension and match the associated MS/MS spectra to (modified) peptide sequences (37). The calibrated inverse reduced ion mobility value (*1/K*_*0*_) determined from the mobility spectrum of each feature can then be converted into ion-nitrogen ^TIMS^CCS_N2_ values using the Mason-Schamp equation (42).

**Figure 1.**
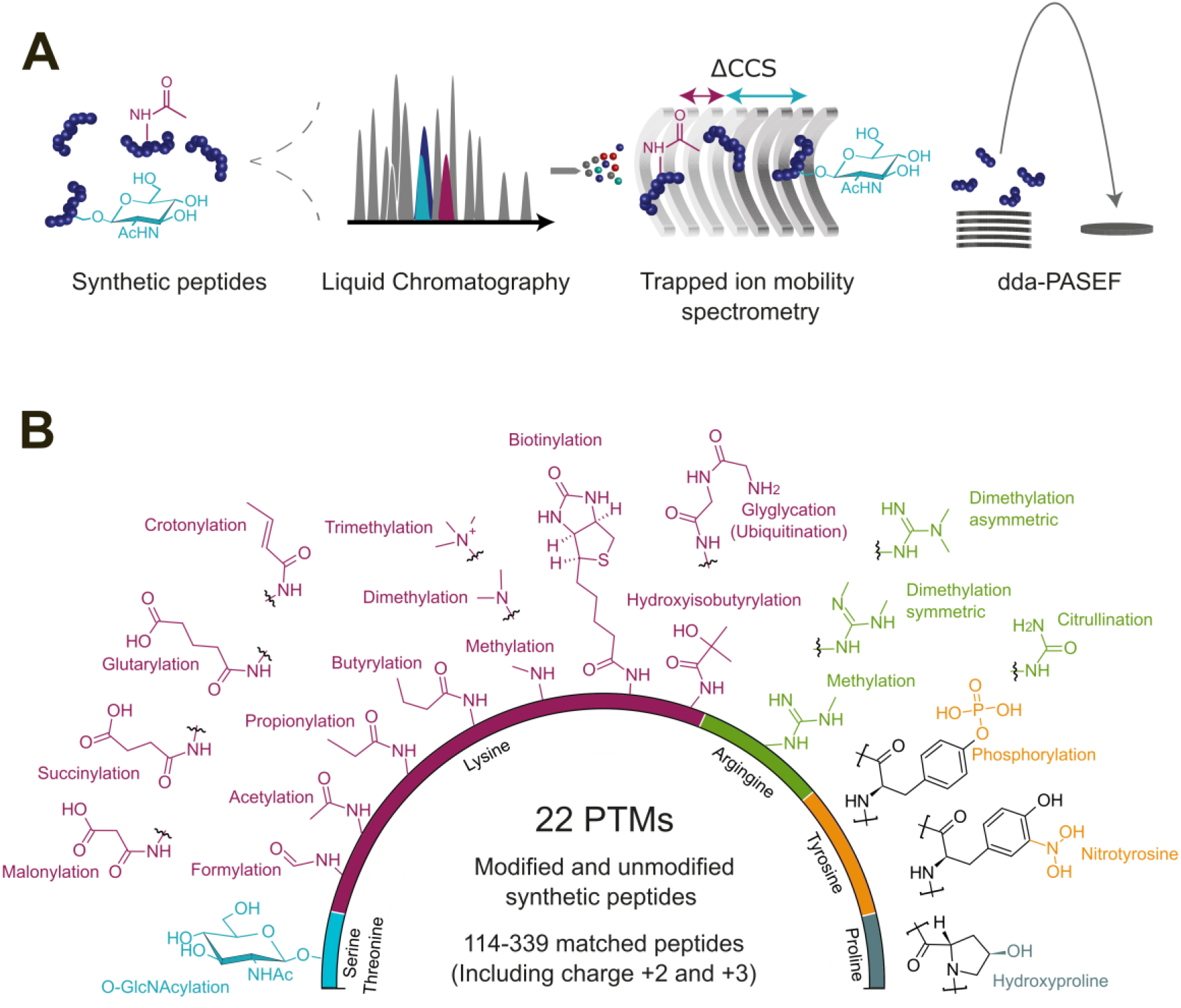
Overview of 22 post-translational modifications (PTMs) analyzed by trapped ion mobility spectrometry. **(A)** Schematic workflow for the analysis of synthetic peptide pools by liquid chromatography and trapped ion mobility - quadrupole time-of-flight mass spectrometry with data-dependent acquisition (dda)-PASEF. **(B)** Dataset summary and chemical structures of the peptide modifications investigated in this study color-coded by their respective modified amino acid.

**Figure 1b** shows an overview of all PTMs in our dataset. The ProteomeTools library contributed matching modified and unmodified tryptic peptides derived from human protein sequences, which were selected for their favorable LC-MS properties and synthesizability as described in more detail in the original publications (34, 35). The library contains four different sets of base peptide sequences that carry N-terminal or internal modifications on one of four amino acids (lysine, arginine, tyrosine and proline). All peptide sequences with the same modification are combined, resulting in 21 pools of modified peptides and four matching pools of unmodified peptides, which we measured in randomized order and in triplicates. In addition, we analyzed a pool of *O*-GlcNAcylated peptides (serine/threonine) and their unmodified counterparts in the same way. Overall, lysine modifications represent the largest group in our dataset, including three homologous series for acylation with aliphatic residues (formylation to butyrylation), carboxylic acid residues (malonylation to glutarylation) and methylation. Further, lysine biotinylation, a particularly bulky modification, is included as well as the GlyGly remnant of ubiquitination and hydroxylated proline. Tyrosine modifications in our dataset are phosphorylation and nitration. The arginine pool adds further subtleties through symmetric and asymmetric di-methylations.

In total, we compiled 84 raw files, which resulted in ∼57,000 peptide spectrum matches mapping to ∼5000 unique combinations of peptide sequence, charge state and modification. Of these, 54% and 43% were detected as doubly and triply charged species, and only 3% in charge state 4. The median Andromeda score was 105 with a very high localization probability close to 1 for all modifications except for O-GlcNAc, which is labile in collision induced dissociation experiments (Supplementary Fig. 1). Plotting the *m/z* vs. CCS distribution of modified and unmodified peptides shows the expected clustering by charge state distributed over an *m/z* range of about 300– 1200 and a CCS range of about 300–700 Å^2^ (Supplementary Fig. 2). For further analysis, we extracted the CCS value of the most abundant feature for each modified peptide sequence, while keeping doubly and triply charged ions separate because of their distinct ion mobility. This yielded approximately 4700 matched pairs of modified and unmodified peptides and about 114 to 339 pairs per modification.

### Precision of TIMS CCS measurements

Experimental ion mobility values depend on the analyte itself as well as the electric field and the nature of the mobility gas. To make them comparable between experiments, TIMS is usually calibrated by a linear regression of known *1/K*_*0*_ values to the elution voltage, resulting in good agreement with conventional drift tube experiments (18). In our previous study, we demonstrated that ^TIMS^CCS values from multiple experiments can be linearly aligned based on overlapping peptide identifications, leading to a remarkable reproducibility over long periods of time and across instruments (13). To enable a similar alignment in our dataset of non-overlapping peptide pools and avoid external calibration before each injection, we here spiked a standard of eleven synthetic iRT peptides into each sample. The peptides eluted evenly distributed over the 2 h chromatographic gradient and were detected as doubly protonated species with CCS values ranging from 320-430 Å^2^. Our analysis revealed a time-dependent shift of their measured CCS values over the course of the experiment (84 LC-MS injections, 10 days), which we attributed predominantly to changes in the ambient air pressure (**Fig. 2a**). A linear alignment to the median CCS values from three reference experiments successfully corrected these drifts (Methods, **Fig. 2a** lower panel). We found that using the median deviation of all detected iRT peptides as a correction factor makes this approach also robust to outliers resulting from peptides with complex ion mobility spectra (e.g. YIAL… in **Fig. 2a**). Across all experiments, the median coefficient of variation of the iRT peptide CCS values was 1.44% before and 0.46% after the alignment.

**Figure 2.**
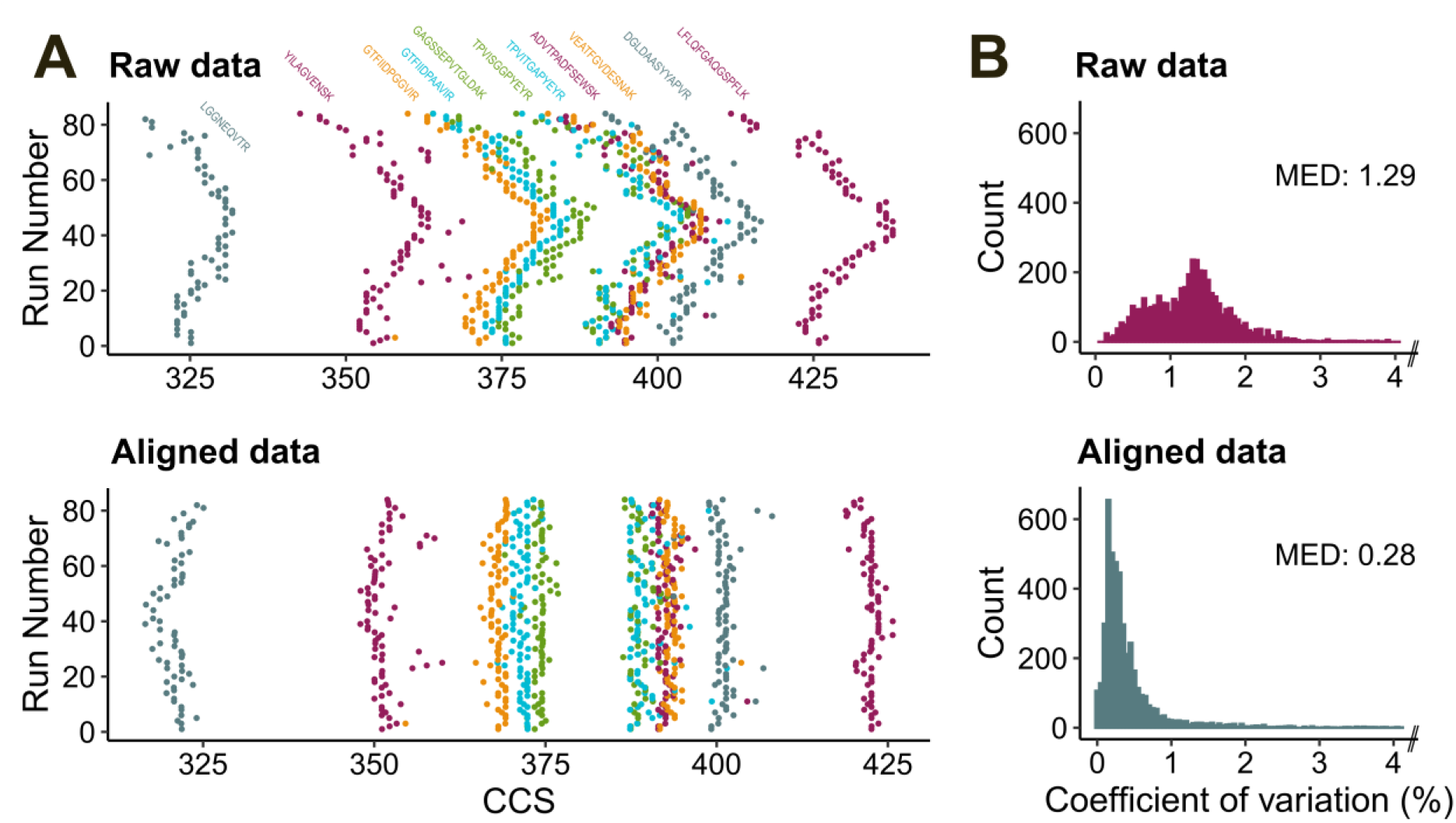
Reproducibility and cross-run alignment of peptide collision cross sections. **(A)** CCS values of eleven reference peptides spiked into 84 consecutive LC-MS experiments. Data are shown in chronological order before (top) and after (below) performing a linear alignment to correct for drifts in the ion mobility measurement (Methods). **(B)** Coefficients of variation of CCS values of (modified) peptides measured in three replicate injections in the full dataset (*top*: raw data, *bottom*: aligned data). N = 4,495, bin-width = 0.05, 1% outliers out of bounds not shown.

Next, we determined the precision of the CCS measurements for all other peptides in our dataset (**Fig. 2b**). Because of the high sequencing rate of dda-PASEF and the relatively low complexity of the synthetic peptide pools, 73% of all peptides were identified in all three replicates. For these, the coefficient of variation was 1.29% before and only 0.28% after linear alignment. This indicates an excellent reproducibility of our TIMS measurements and is in line with previous reports on this instrument platform (5, 13).

### Global view on CCS values of modified peptides

The fact that charge is a major determinant of ion mobility prompted us to investigate the occurrence of different charge states in our dataset. **Figure 3a** provides an overview of predominant charge states as well as the relative abundance fraction of charge states for all peptides. Tryptic peptides generally take up two to four protons in the electrospray process, depending on the length of the amino acid sequence and the number of basic residues. PTMs can alter the gas phase basicity of the modified amino acid and hence the charge state distribution. This effect was most striking for lysine modifications, as doubly and triply charged species appeared in roughly equal abundance for the unmodified peptides, whereas the various acylations shifted the charge distribution almost completely to charge 2. By contrast, lysine methylations as well as the GlyGly residue retain basic properties at the lysine site and thus showed only little effect on the relative abundance distribution, but rather tipped the predominant charge state to 3. We observed similar trends for arginine methylation and, as expected, citrullination reduced the charge state. Hydroxylation of proline, O-GlcNAcylation of serine and threonine as well as nitration and phosphorylation of tyrosine did not alter the charge state distribution as compared with their unmodified counterparts. Overall, these results are in agreement with the preceding analysis of the 21 PTM library on a different instrument platform (35).

**Figure 3.**
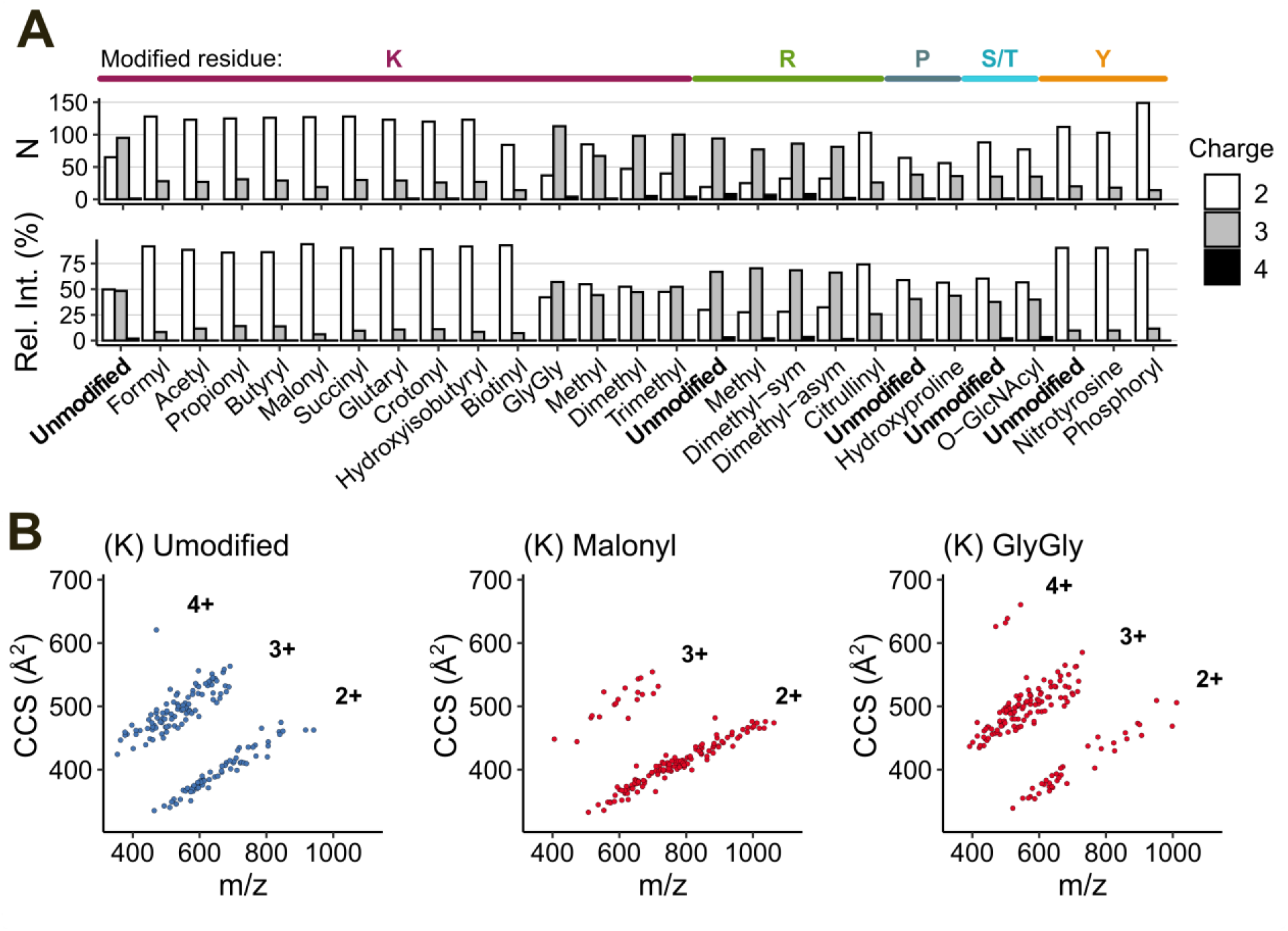
Analysis of charge state and collision cross section (CCS) distribution for different PTMs. (**A**) Number of identified peptides per modification in their predominant charge state (top). Relative abundance of different charge states for all identified peptides (bottom). For R and K, only unmodified peptide sequences with at least one internal R or K are shown. **(B)** Peptide *m/z* vs. CCS distribution of selected modifications from our dataset of 22 PTMs. The scatter plots show peptides in their predominant charge state.

To illustrate the global effect of charge state alterations on the ion mobility of modified peptides, we plotted them in their predominant charge state in the *m/z* vs. CCS space (**Fig. 3b**). The unmodified peptides of the lysine pool (left panel) are not strictly tryptic peptides because of their internal lysine. Nevertheless, they followed the well-characterized distribution of ion mobility and charge state occupancy. Malonylation, as an example for the group of acylations, thinned the population of charge 3 species and led to a more dense population of charge 2 peptides. Conversely, GlyGlycation caused a distinct shift in the opposite direction and predominantly populated the area of triply charged species. Thus, simple charge state alterations can already contribute to the separation of peptides and modifications in the ion mobility dimension.

### Pairwise comparison of modified and unmodified peptides

A distinct feature of our data set is the large number of matching pairs of modified and unmodified peptide sequences. Thus, having determined the position of modified peptide populations in the CCS vs. *m/z* space, we next performed a pairwise analysis of the respective counterparts. Intuitively, one could expect increasing CCS values throughout, as all modifications in our dataset introduce an additional functional group to the peptide (with the only exception of citrullination). However, we observed relative differences (ΔCCS = (CCS_modified_ – CCS_unmodified_)/CCS_unmodified_) ranging all the way from about -10% to +10%, depending on the type of modification, the modified amino acid as well as the charge state (**Fig. 4a**). As a proxy for the experimental precision in the data set, we indicate the +/- three-fold coefficient variation (0.84%) interval in the boxplot. The median ΔCCS shifts of specific modifications ranged from almost no difference (0.7% for tyrosine nitration), to slightly negative values (−1.1% for lysine formylation) and clear positive shifts for bulky modifications such as biotinylation (4.3%) and O-GlcNAcylation (4.5%).

**Figure 4.**
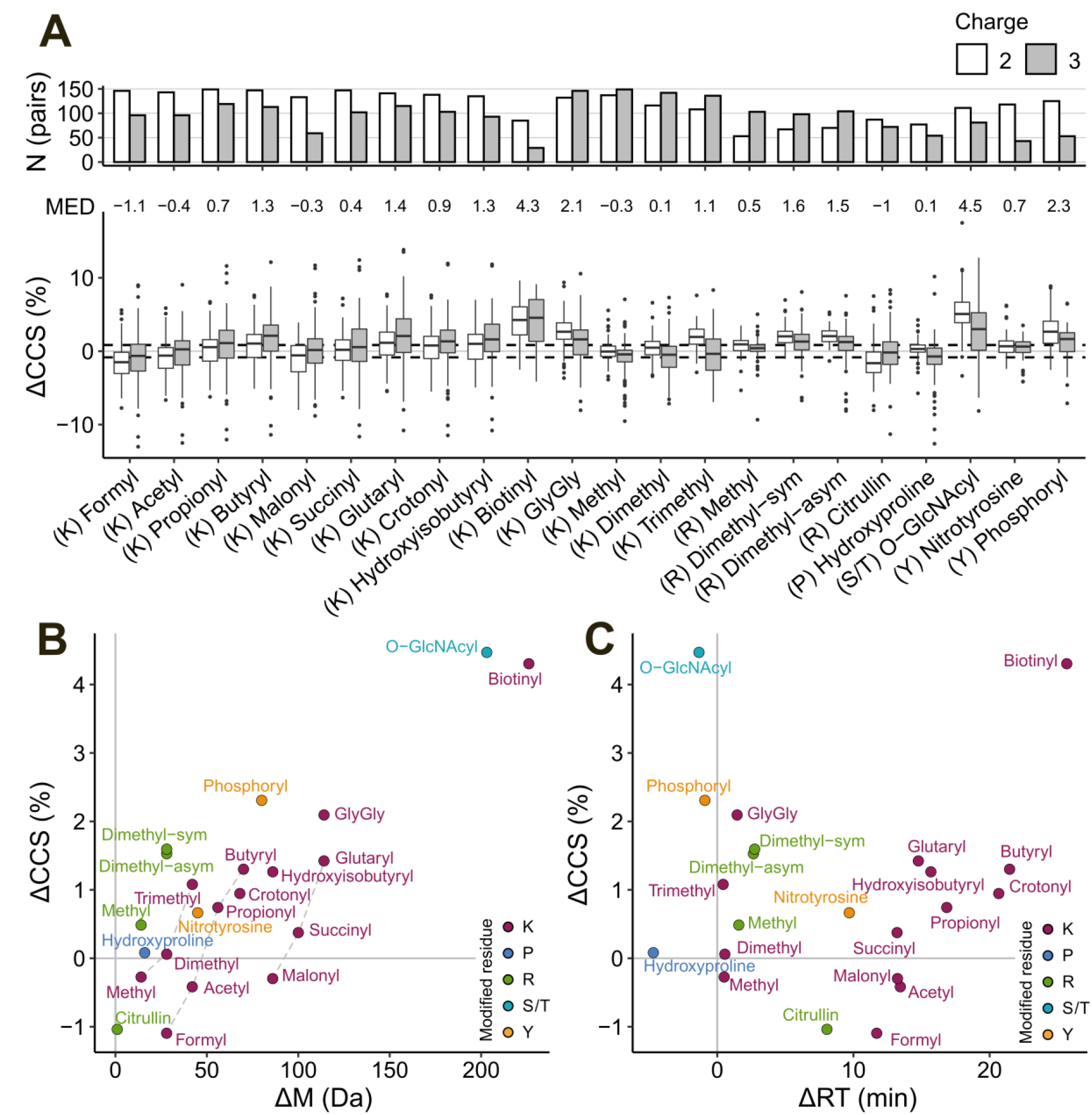
Pairwise comparison of modified and unmodified peptide collision cross section (CCS). **(A)** Relative difference of CCS values of matching pairs, separated by charge state (ΔCCS). The number of pairs is plotted on the top axis. The median relative deviation (MED) indicates the combined value for charge 2 and 3. Dashed lines indicate the 3 x median coefficient of variation in the full data set as a proxy for experimental precision. Boxplot elements: Interquartile range within boxes; median indicated by horizontal line; whiskers spanning 1.5 x interquartile range. **(B)** Median ΔCCS values of all investigated modifications as a function of the molecular weight change (ΔM). Dashed lines visualize homologous series. **(C)** Same as B, but as a function of the observed shift in retention time (ΔRT).

To delve deeper into factors that determine CCS values of modified peptides, it is insightful to first consider the two different homologous series in the subset of lysine modifications: acylations with mono- (formyl to butyryl) and di-carboxylic acids (malonyl to glutaryl). Both series show a nearly linear increase in CCS with longer acyl chains, consistent for both charge 2 and 3. This is in line with the intuition of increasing bulkiness with increasing chain length. To our surprise, and despite the fact that both series differ by one carboxylic acid, the median shifts in CCS of, for example, acetyl and malonyl were almost identical (−0.4% and -0.3%). Moreover, they were similar to that of lysine methylation (−0.3%), an even smaller moiety. A possible explanation for this observation are the different chemical properties of the modifications. While methylation tends to increase the basicity at the modified lysine, the acetyl group is electron withdrawing and malonylation adds an acidic functionality. Peptide conformations in the gas phase can be partially explained by Coulomb interactions and intramolecular charge solvation (11). Analyzing a larger data set of peptide CCS values, Chang *et al*. concluded that internal acidic residues facilitate more compact conformations and, conversely, internal basic residues result in more extended conformations with larger cross sections. Our data is in line with this model and suggests that the carboxyl residue of malonyl compensates for its increased steric bulkiness.

The correlation between peptide mass and cross section is well studied. Interestingly, we observed a lower correlation between CCS and *m/z* for modified (charge 2: r^2^ = 0.90, charge 3: r^2^ = 0.67) than for unmodified peptides in our dataset (charge 2: r^2^ = 0.93, charge 3: r^2^ = 0.75). To investigate this in more detail, we plotted the median ΔCCS values as a function of the modification’s molecular weight (**Fig. 4b**). This analysis reproduced the large effect size for O-GlcNAcylation and biotinylation and showed an overall correlation of r^2^ = 0.66. However, when excluding the latter from the analysis, r^2^ dropped to only 0.25, indicating that mass alone is a poor predictor of ΔCCS values for this group of chemically diverse modifications. In line with the results above, the homologous lysine modification series appear on parallel lines in this plot (dashed grey lines). Modifications with similar ΔCCS but different molecular weight such as the methyl-acetyl-malonyl example above (7, 21 and 43 Da) appear on thought horizontal lines. Vertical lines, on the other hand, show modifications with similar molecular weight but different ΔCCS values. As the example of lysine- and arginine-dimethylation shows, this effect is not limited to the chemistry of the modification itself, but can also depend on the modified amino acid. Performing a similar correlation analysis of ΔCCS and changes in the chromatographic retention time (**Fig. 4b**), revealed some degree of orthogonality between LC and ion mobility. O-GlcNAcylation, phosphorylation and GlyGlycation, for example, only slightly altered the retention time, while resulting in relatively large CCS shifts.

Although our analysis highlighted conceivable trends for each modification in our data, we also noted a relatively high variance within the modification groups (Fig. 4a). In most cases, the inter-quartile range of the pairwise analysis was several-fold larger than the precision of our experiments. Furthermore, some modifications showed less variance compared to others and the variance in charge 3 species was generally higher. This hints towards sequence-dependent effects within the peptide pools that modulate the effect of modifications on peptide cross sections.

### Resolving sequence-dependent CCS determinants

To dissect the high variance within the modification groups, we next resolved our data by peptide sequences. This is possible because all peptides from one amino acid pool have the same unmodified base sequence. **Figure 5a** shows absolute ΔCCS values of all doubly charged peptides in the subset of lysine modifications color-coded by their respective base peptide sequence. Strikingly, the rank order of ΔCCS values remained largely unchanged throughout the homologous series of acyl modifications. In other words, while a particular modification could alter a peptide’s cross section from +10 Å^2^ to -25 Å^2^ (formylation) depending on its amino acid sequence, elongating the modification, e.g. from formyl to butyryl, resulted in consistent increments across all peptide sequences. This observation also applied to triply charged peptide ions (Suppl. Fig 3) and only few peptides deviated from this trend. Even for the chemically rather distant biotinylation, we observed a similar behavior, although more peptides swapped positions, in particular those with larger ΔCCS values. In contrast, when extending the line plot to lysine methylations and GlyGly, the trend was interrupted. However, the latter modifications are also distinct from the former acyl-type modifications in terms of charge state distribution (**Fig. 3**). This suggests that the observed grouping is driven by intramolecular charge localization and solvation. Similarly, we found that for individual sequences, the absolute ΔCCS as well the shift relative to other sequences can be discordant for charge states (**Fig. 5b**).

**Figure 5.**
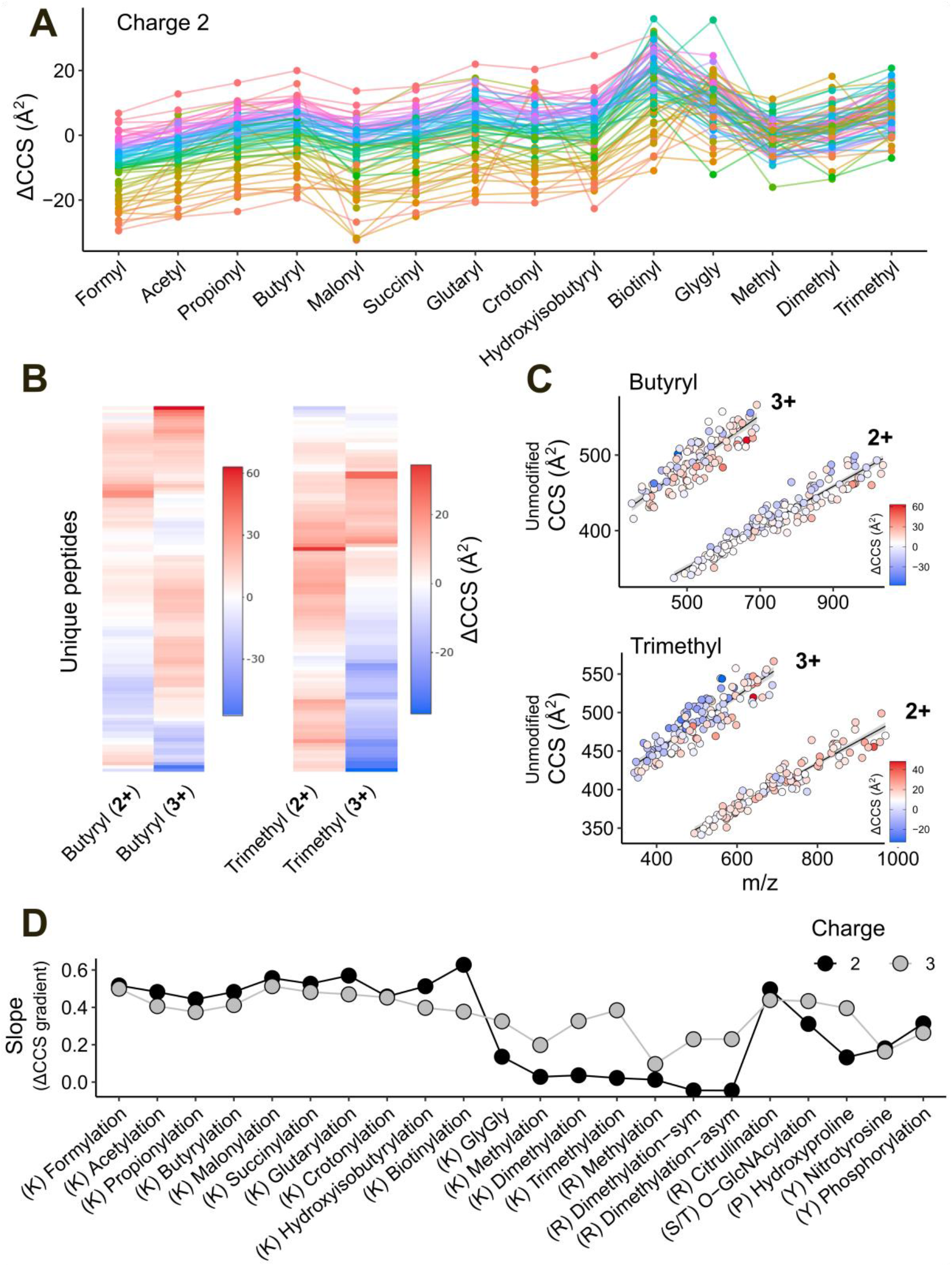
Sequence-dependent fine structure of PTM-induced CCS shifts. **(A)** ΔCCS values of individual peptide sequences for lysine-modifications at charge 2. Only peptides which were detected for all shown modifications are included (N = 73). **(B)** Heat maps of ΔCCS values comparing charge 2 and 3 for both butyrylation (N = 108) and trimethylation (N = 101). **(C)** Peptide *m/z* vs. CCS distribution of unmodified reference peptides for butyrylation (N = 260) and trimethylation (N = 244), overlaid with each peptides PTM-induced CCS shift. Linear trend lines are fitted to both charge states. **(D)** ΔCCS gradient within ion clouds for each investigated PTM, separated by charge. Each peptides residual to a linear regression, as depicted in panel C, was plotted against its ΔCCS value. The slope of a subsequent linear regression represents a ΔCCS gradient within the respective ion clouds (panel C).

These results raised the question of whether there are commonalities in peptide sequences that undergo similar changes in their gas phase structure upon modification. To this end, we plotted the *m/z* vs. CCS distribution of the unmodified peptides and overlaid the corresponding ΔCCS values for different modifications. As a visual aid, we divided them into extended (larger CCS values) and compact structures (smaller CCS values) by fitting a linear model to each charge state. **Figure 5d** shows lysine butyrylation and trimethylation as examples. For butyrylation, close inspection of the doubly and triply charged ion populations revealed a tendency towards negative ΔCCS values for extended structures and vice versa. We also observed a compaction of extended structures for triply charged trimethylated peptides, indicating that the internally localized charge destabilizes extended conformations. This sequence-dependency of trimethylation was less pronounced for doubly charged species, in line with their narrower ΔCCS distribution in Fig. 4a.

Taken our results together, we hypothesized that the gas-phase conformation of the unmodified peptide ion modulates the relative effect of specific modifications. To test this on all our data, we plotted the residuals of the linear fit (indicating whether the unmodified peptide adapts a more compact or extended structure) as a function of the corresponding ΔCCS value for each modification (Suppl. Figs. 4 and 5). The slope of the resulting linear trend lines can be interpreted as ‘ΔCCS gradients’ within the CCS vs. *m/z* ion populations (**Fig. 4d**). Indeed, all lysine acylations and citrullination followed the trend described for butyrylation, while GlyGly and methylated peptides resembled trimethylation. From this, we concluded that the ΔCCS value associated with a modification is indeed dependent on the ‘starting point’ of the unmodified peptide and hence on the amino acid sequence, rather than a fixed increment determined by its chemical composition.

## Conclusions

Recent advances in the application of ion mobility spectrometry to MS-based proteomics also promise new opportunities for the proteome-wide characterization of post-translational modifications. In particular, the combination of TIMS and PASEF has enabled the precise measurement of CCS values on the scale of hundreds of thousands to more than a million data points and contributed to a better understanding of sequence- and position-dependent determinants of peptide cross sections (13, 32, 43). To extend this work beyond unmodified peptides, here we used synthetic peptide libraries with known ground truth to characterize the effect of 22 different PTMs on peptide CCS values.

Our study provides data on 114 to 339 matched modified and unmodified peptides per modification with a precision <1% after linear alignment. This is on par with previous studies on the same instrument platform (5, 13) and enabled us to investigate different layers of modification-specific effects on peptide CCS values. On a global level, we observed major shifts in the *m/z* vs. ion mobility distribution for modified peptides, which we attributed to changes in their predominant charge state. In proteomics practice, such effects can be important, for example, to optimize the precursor selection scheme in dia-PASEF experiments (21, 22) or to bias data-dependent acquisition towards modified peptides (24, 44).

Turning to pairwise comparisons of modified peptides and their unmodified counterparts, we observed median ΔCCS values in a range of -1.1% to 4.8%. Surprisingly, despite the correlation between ion mass and mobility, the modification mass alone proved to be a poor predictor of ΔCCS values for most modifications in our data set. In parts, we could rationalize these observations by the counteracting effects of increased bulkiness on the one hand and intramolecular charge solvation on the other hand. In addition, our data revealed substantial sequence-dependent effects on the cross section of modified peptides. This is in line with another recent study focused on phosphorylated peptides (23). All in all, our study adds to the increasing body of work indicating that peptide cross sections are determined by the amino acid composition (45, 46) as well as their linear sequence (13, 32, 47).

Accurate predictions of peptide properties such as retention time, MS/MS spectra as well as CCS values are increasingly used in MS-based proteomics (48, 49). In this context, synthetic peptides can provide important training data as they have a known ground truth (34). This applies in particular to modified peptides, which are not always readily accessible from biological sources via efficient and affordable enrichment protocols. We envision that our high-quality data set fills this gap and helps to extend CCS prediction algorithms to various post-translational modifications, for example via transfer learning (33).

## Acknowledgements

This work was partially supported by the Federal Ministry of Education and Research and the Thuringian Ministry for Economic Affairs, Science and a Digital Society through the Joint Federal Government-Länder Tenure-Track Programme, by the Free state of Thuringia and the European Union via the “Innovationszentrum für Thüringer Medizin technik-Lösungen” (ThIMEDOP; #2018 IZN 002), by the Center for Interdisciplinary Clinical Research (IZKF-Jena) and by the German Research Foundation through the Research Training Group ‘ProMoAge’ (RTG2155). We thank our colleagues at the Jena University Hospital for fruitful discussions and technical support, in particular F. Schneidmadel and C. Tschernjawski. The 21 PTM peptide library was synthesized as part of the ProteomeTools project and kindly gifted by the Küster laboratory (34, 35). We acknowledge M. Wilhelm for providing details about the peptide library and valuable discussions.

