## Supplementary Information for "Peptide collision cross sections of 22 post-translational modifications"

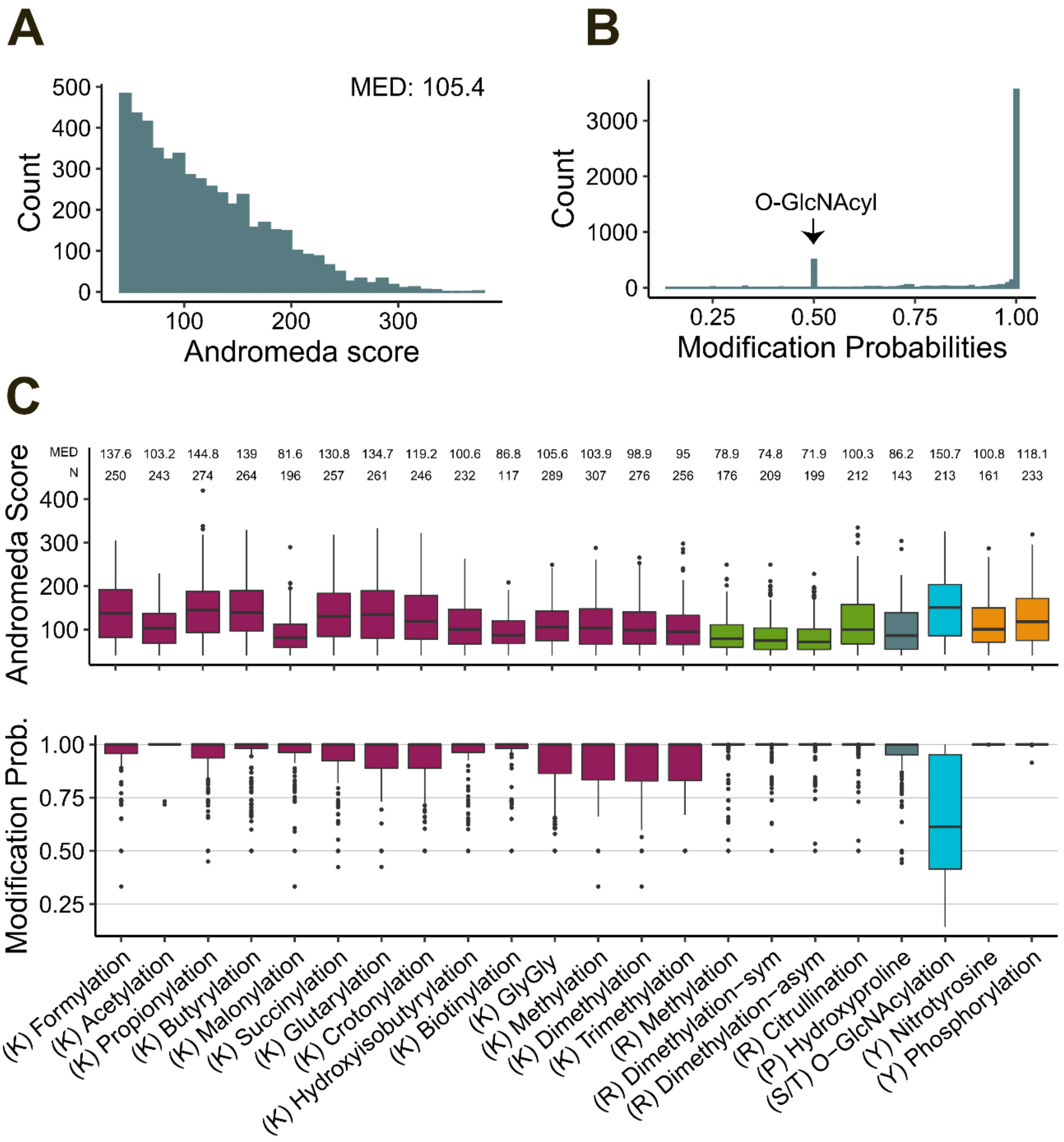


**Supplementary Figure 1. (A)** Andromeda score distribution of unique (highest intensity) combinations of sequence, charge state and modification (n = 5014, bin width = 10). **(B)** Same as A, but for modification localization probability (bin width = 0.01). **(C)** Modification specific boxplot of Andromeda score distributions (top) and modification localization probability (bottom). Boxplot elements: Interquartile range within boxes; median indicated by horizontal line; whiskers spanning 1.5 x interquartile range.

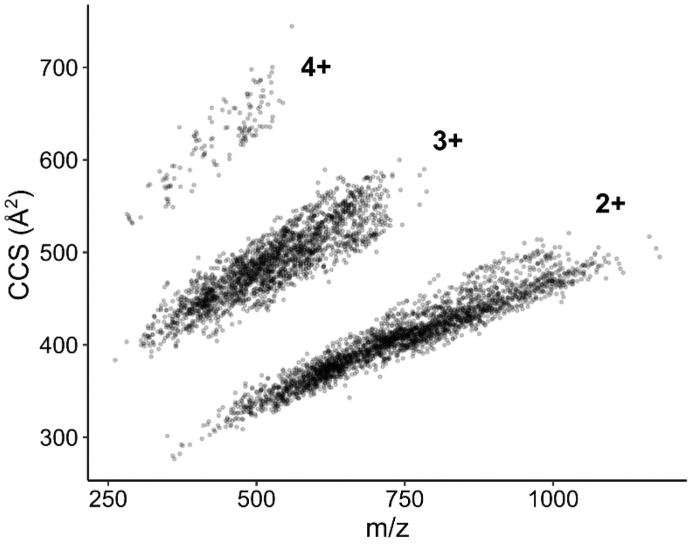


**Supplementary Figure 2.** Distribution of unique combinations of sequence, charge state and modification in the *m/z* vs. CCS space (n = 5014).

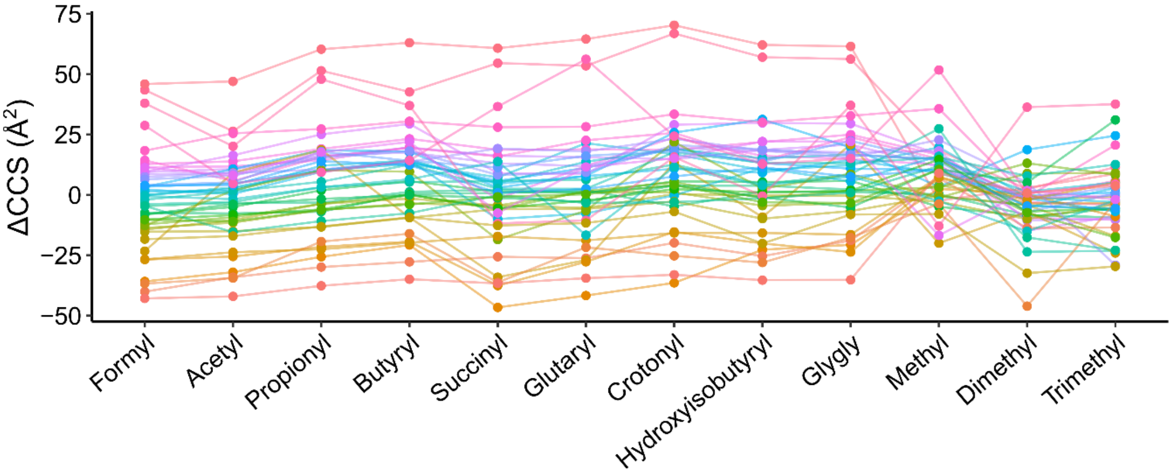


**Supplementary Figure 3.** ΔCCS values of individual peptide sequences for acyl type modifications on lysine at charge 3. Malonylation and biotinylation were excluded due to their low intersection (n = 45).

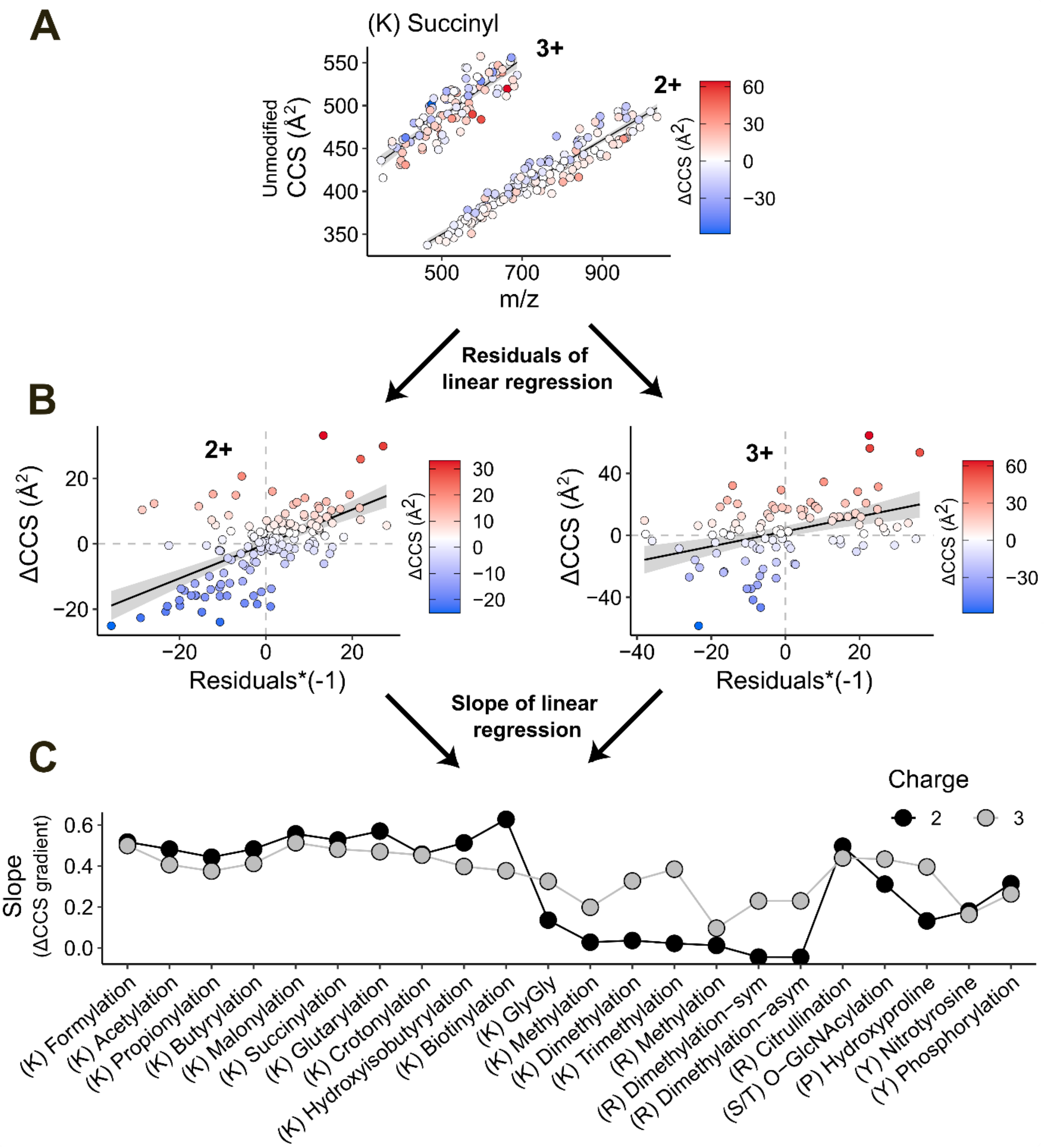


**Supplementary Figure 4. Schematic workflow for the analysis of ΔCCS gradients.** (A) Peptide *m/z* vs. CCS distribution of unmodified peptides color-coded by ΔCCS values for succinylation. Linear regression lines are fitted to both charge states. (B) ΔCCS vs. residuals of the linear regression from A. To observe trends, a liner regression is fitted to this graph as well. We interpreted the resulting slope as a 'ΔCCS gradient' within the ion populations shown in panel A. (C) Overview of ΔCCS gradients as described in panel B for all investigated modifications.

**Supplementary Table 1.** Parameters for MaxQuant modification search.

| Modification | Mass (Da) | Residue | Composition |
|---|---|---|---|
| Acetylation | 42.011 | K | H(2) C(2) O |
| Biotinylation | 226.078 | K | H(14) C(10) N(2) O(2) S |
| Butyrylation | 70.042 | K | H(6) C(4) O |
| Crotonylation | 68.026 | K | H(4) C(4) O |
| Dimethylation | 28.031 | K | H(4) C(2) |
| Formylation | 27.995 | K | C O |
| Glutarylation | 114.032 | K | H(6) C(5) O(3) |
| GlyGly | 114.043 | K | H(6) C(4) N(2) O(2) |
| Hydroxyisobutyrylation | 86.037 | K | H(6) C(4) O(2) |
| Malonylation | 86.000 | K | H(2) C(3) O(3) |
| Methylation | 14.016 | K | H(2) C |
| Propionylation | 56.026 | K | H(4) C(3) O |
| Succinylation | 100.016 | K | H(4) C(4) O(3) |
| Trimethylation | 42.047 | K | H(6) C(3) |
| Hydroxyproline | 15.990 | P | O |
| Citrullination | 0.984 | R | H(-1) N(-1) O |
| Dimethylation-asymmetric | 28.031 | R | H(4) C(2) |
| Dimethylation-symmetric | 28.031 | R | H(4) C(2) |
| Methylation | 14.016 | R | H(2) C |
| O-GlcNAcylation | 203.079 | S/T | H(13) C(8) N O(5) |
| Nitration | 44.985 | Y | H(-1) N O(2) |
| Phosphorylation | 79.966 | Y | H O(3) P |

**Supplementary Table 2.** Median collision cross section (CCS), retention time (RT) and mass (m) of three reference measurement of 11 iRT peptides.

| Sequence | CCS (Å$^2$) | RT (min) | m (Da) |
|---|---|---|---|
| LGGNEQVTR | 321.83 | 20.09 | 974.51 |
| YILAGVENSK | 351.09 | 49.81 | 1094.60 |
| GTFIIDPGGVIR | 369.18 | 85.34 | 1245.71 |
| GTFIIDPAAVIR | 372.37 | 96.33 | 1273.74 |
| GAGSSEPVTGLDAK | 374.50 | 38.08 | 1289.65 |
| TPVITGAPYEYR | 387.35 | 59.67 | 1367.71 |
| TPVISGGPYEYR | 387.43 | 55.53 | 1339.68 |
| ADVTPADFSEWSK | 391.47 | 70.74 | 1453.67 |
| VEATFGVDESNAK | 394.96 | 44.78 | 1367.66 |
| DGLDAASYYAPVR | 400.30 | 65.95 | 1398.68 |
| LFLQFGAQGSPFLK | 422.63 | 99.84 | 1553.86 |
