## Supplementary Figure 5 for "Peptide collision cross sections of 22 post-translational modifications"

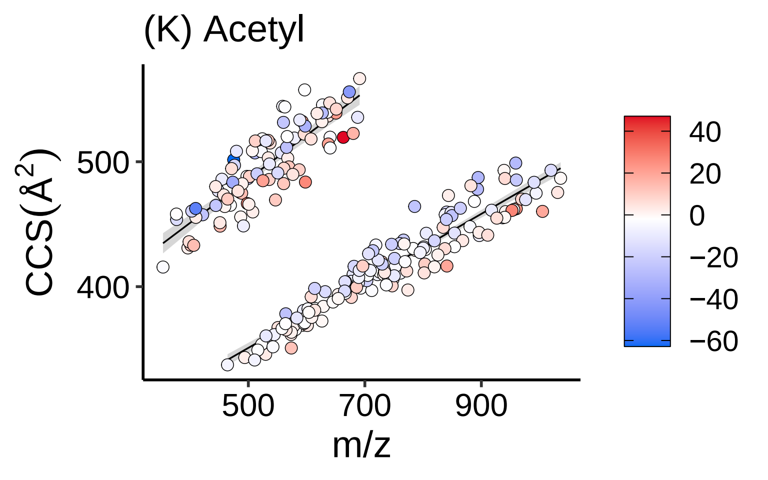

(K) Acetyl
CCS(Å²)
500
400
500
700
900
m/z
40
20
0
−20
−40
−60

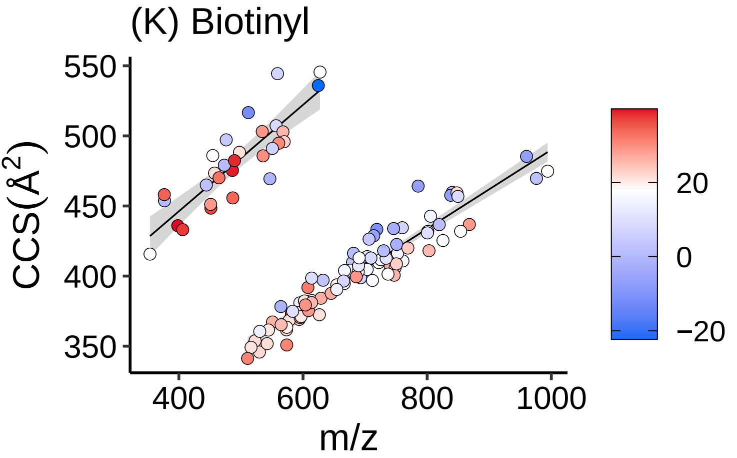

(K) Biotinyl
CCS(Å$^2$)
550
500
450
400
350
400
600
800
1000
m/z
20
0
−20

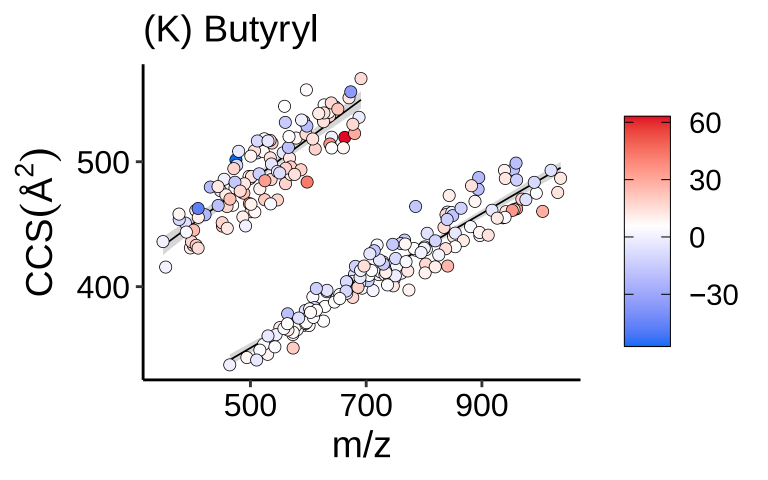

(K) Butyryl
CCS(Å$^2$)
500
400
500
700
900
m/z
60
30
0
−30

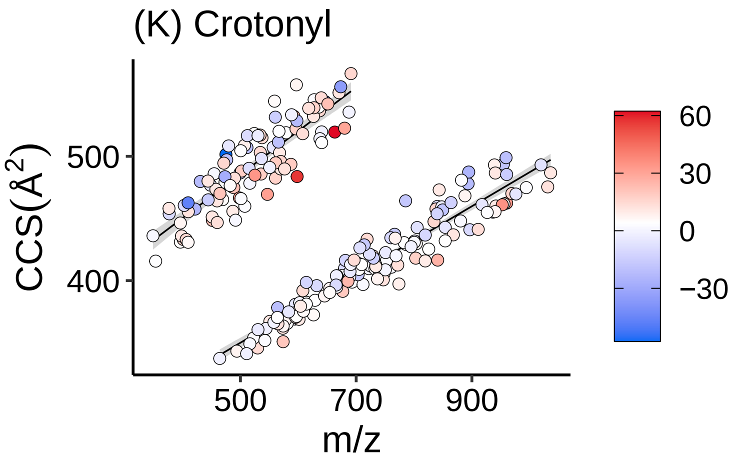
(K) Crotonyl
CCS(Å$^2$)
500
400
500
700
900
m/z
60
30
0
−30

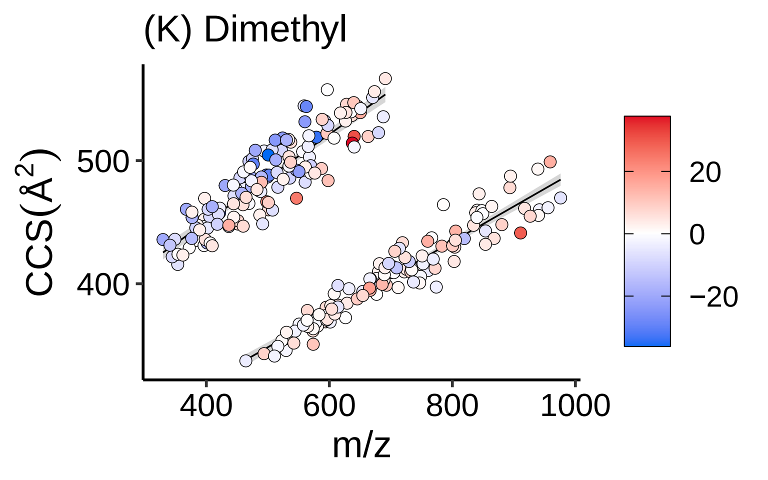

(K) Dimethyl
CCS(Å$^2$)
500
400
400
600
800
1000
m/z
20
0
−20

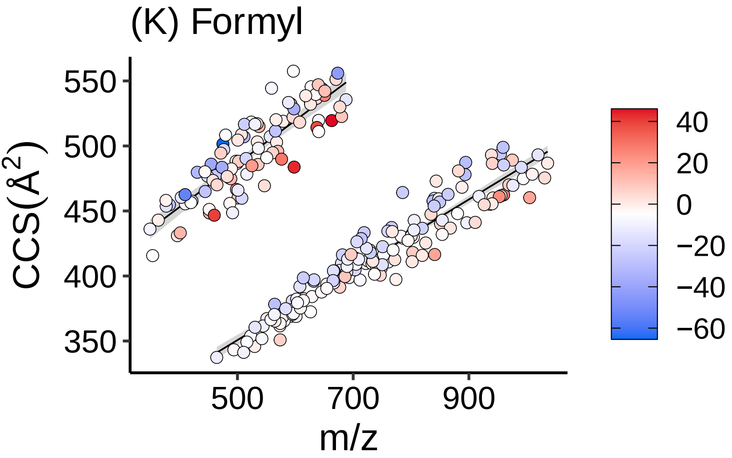
(K) Formyl
550
500
450
400
350
CCS(Å$^2$)
500
700
900
m/z
40
20
0
−20
−40
−60

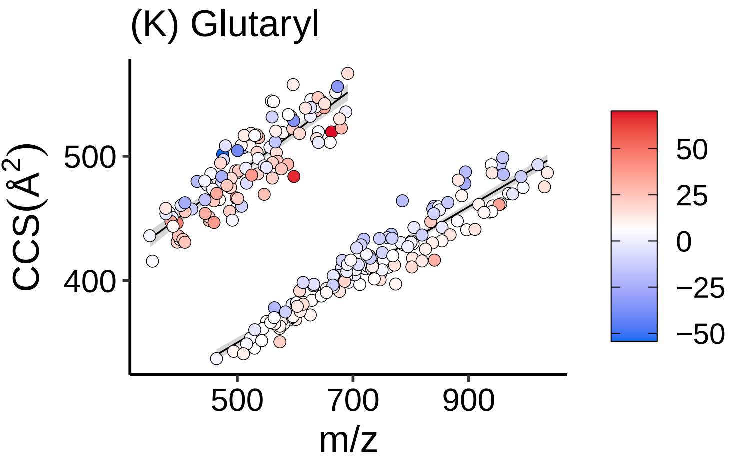
(K) Glutaryl
CCS(Å²)
500
400
m/z
500
700
900
50
25
0
−25
−50

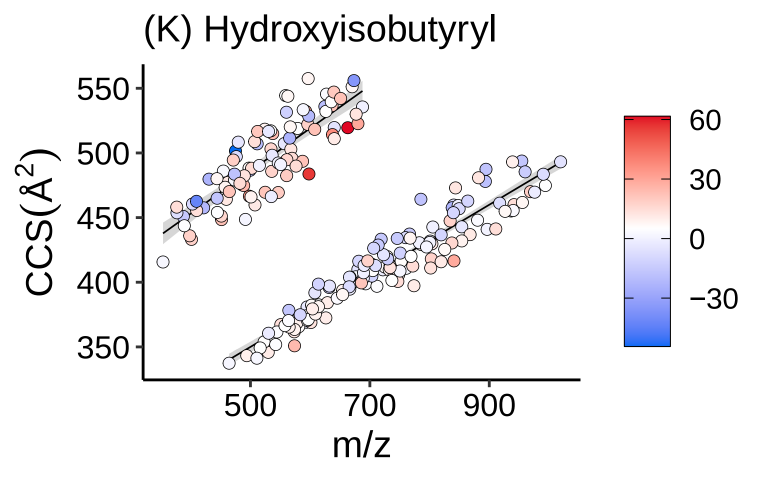

(K) Hydroxyisobutyryl
550
500
450
400
350
CCS(Å$^2$)
500
700
900
m/z
60
30
0
−30

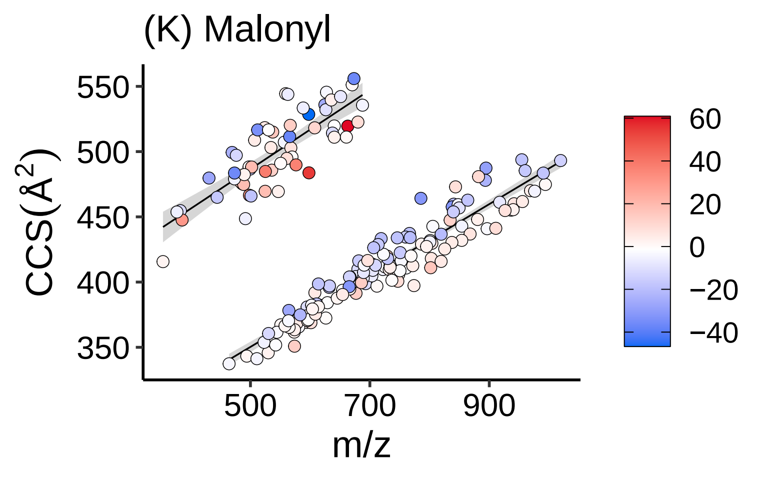

(K) Malonyl
CCS(Å²)
m/z
550
500
450
400
350
500
700
900
60
40
20
0
−20
−40

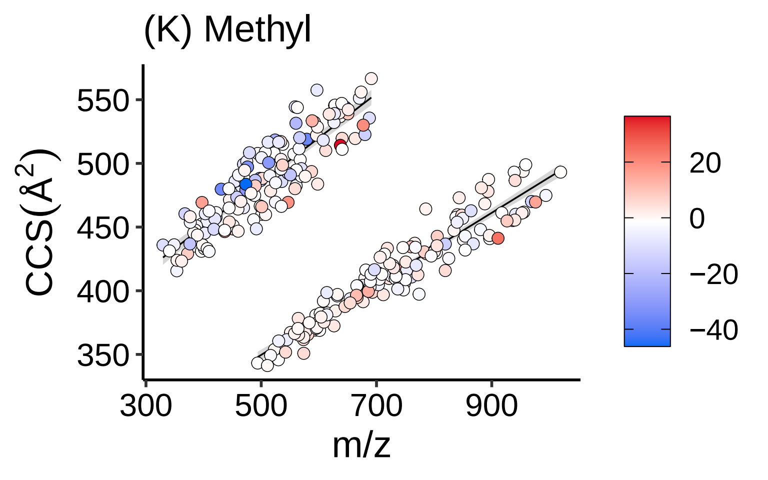
(K) Methyl
CCS(Å$^2$)
550
500
450
400
350
300
500
700
900
m/z
20
0
−20
−40

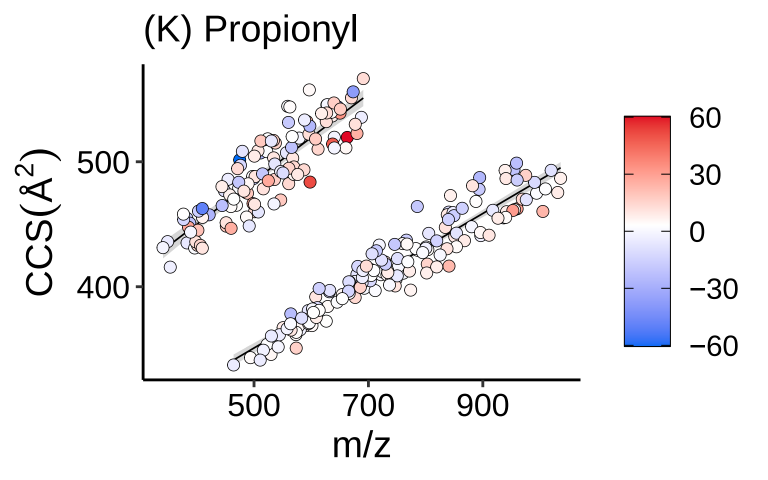

(K) Propionyl
CCS(Å$^2$)
500
400
500
700
900
m/z
60
30
0
−30
−60

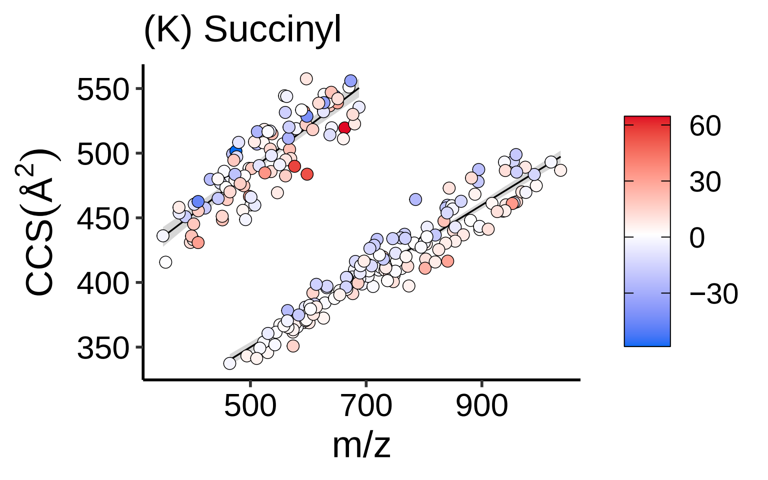
(K) Succinyl
CCS(Å$^2$)
550
500
450
400
350
500
700
900
m/z
60
30
0
−30

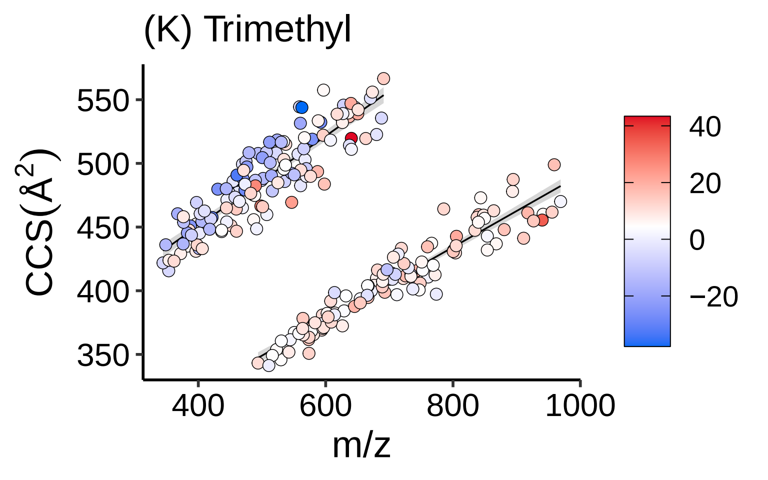

(K) Trimethyl
CCS(Å²)
550
500
450
400
350
400
600
800
1000
m/z
40
20
0
−20

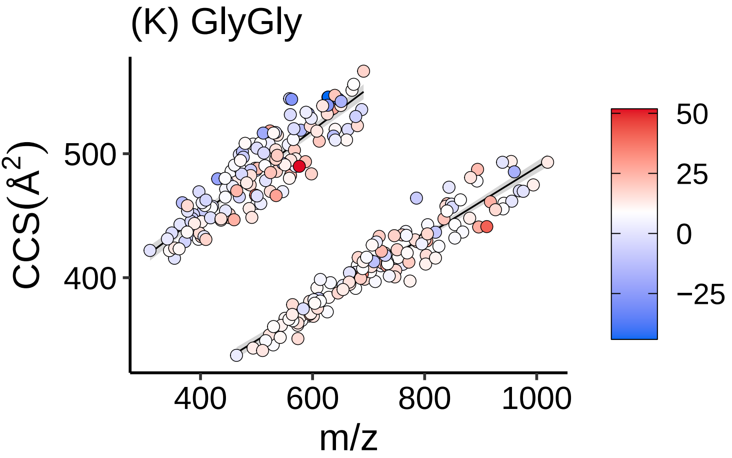
(K) GlyGly
CCS(Å$^2$)
500
400
400
600
800
1000
m/z
50
25
0
−25

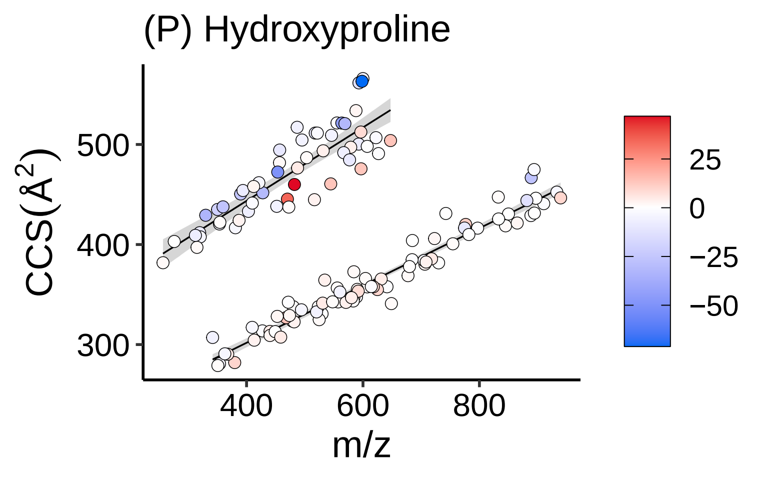
(P) Hydroxyproline
CCS(Å$^2$)
500
400
300
400
600
800
m/z
25
0
−25
−50

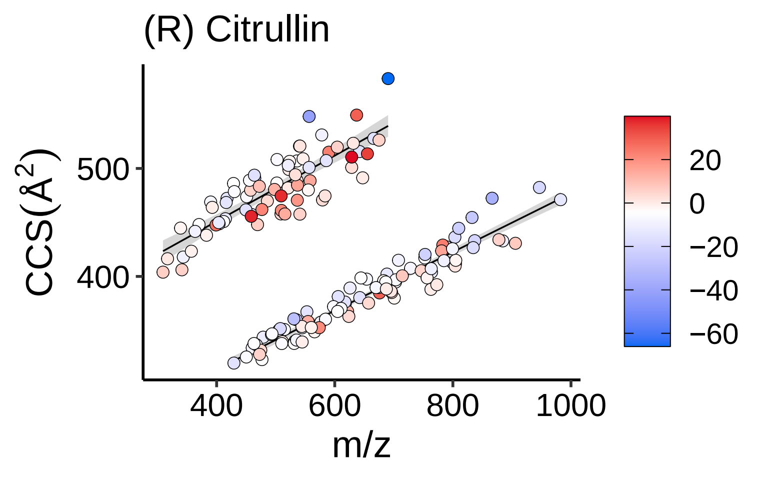
(R) Citrullin
CCS(Å$^2$)
500
400
m/z
400
600
800
1000
20
0
−20
−40
−60

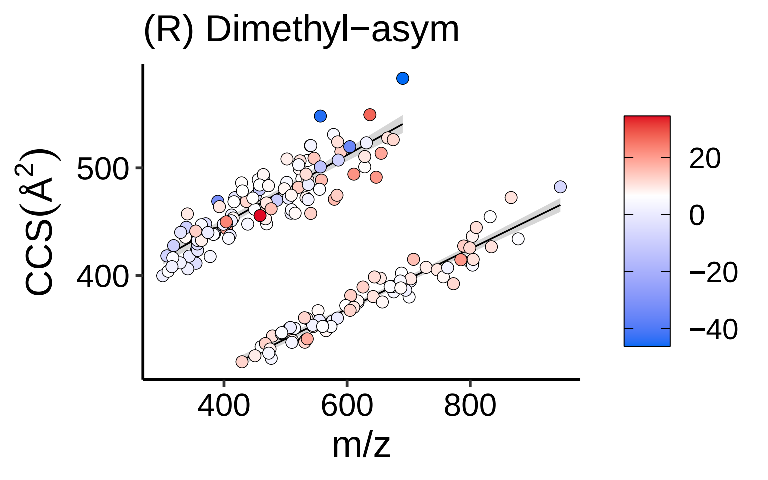
(R) Dimethyl−asym
CCS(Å$^2$)
500
400
400
600
800
m/z
20
0
−20
−40

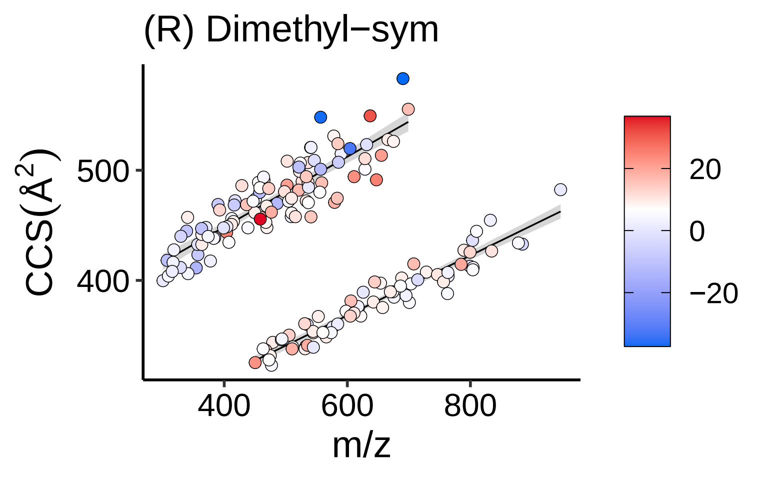

(R) Dimethyl−sym
CCS(Å$^2$)
500
400
400
600
800
m/z
20
0
−20

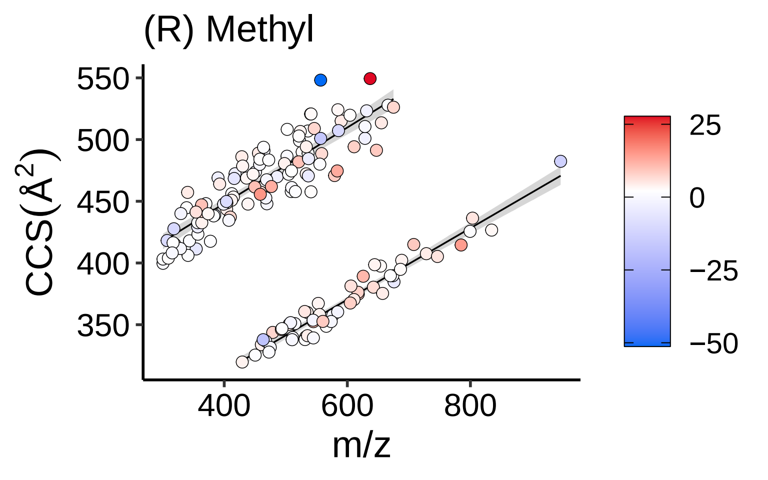
(R) Methyl
CCS(Å$^2$)
550
500
450
400
350
m/z
400
600
800
25
0
−25
−50

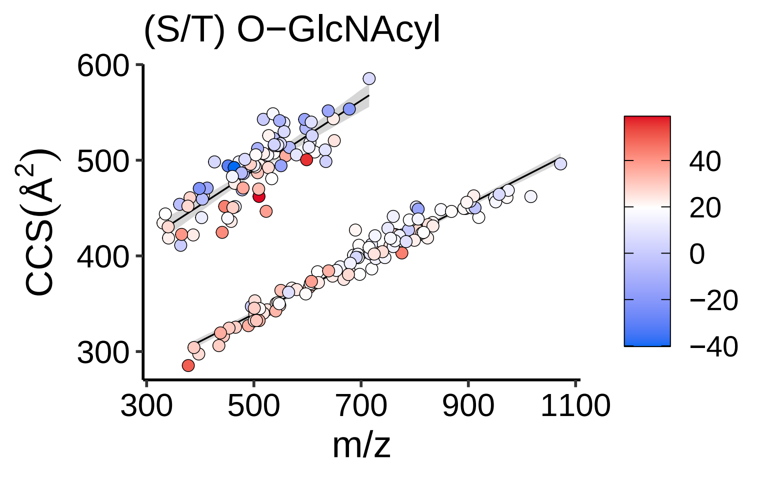

(S/T) O−GlcNAcyl
CCS(Å²)
600
500
400
300
300
500
700
900
1100
m/z
40
20
0
−20
−40

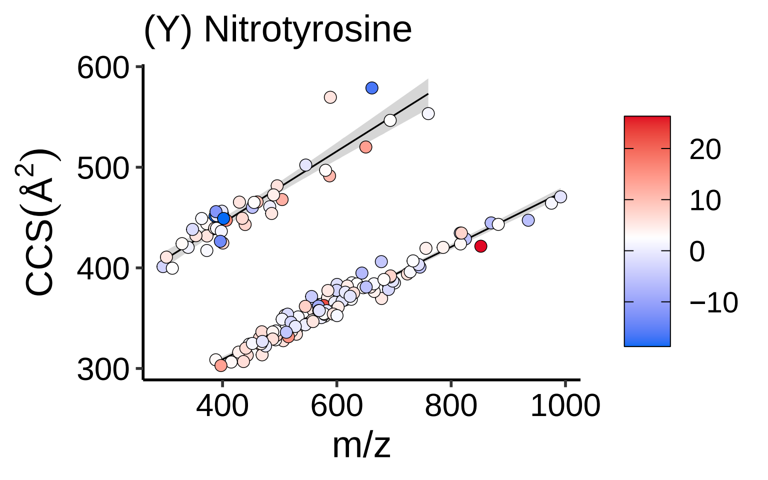

(Y) Nitrotyrosine
600
500
400
300
CCS(Å$^2$)
400
600
800
1000
m/z
20
10
0
−10

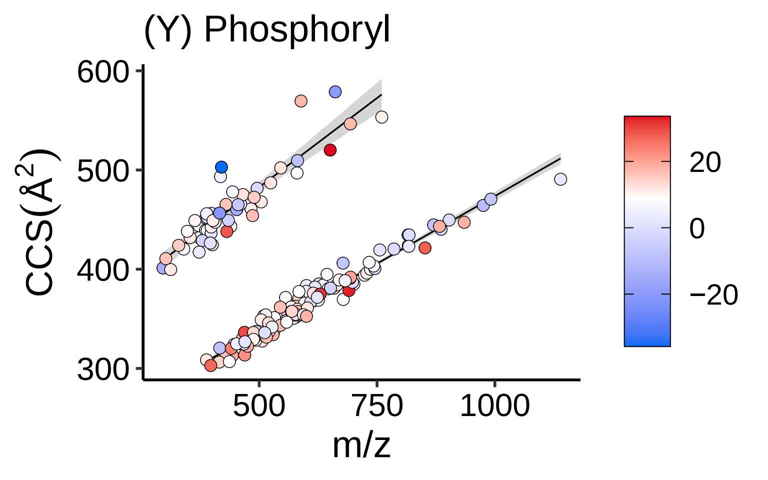

(Y) Phosphoryl
600
500
400
300
CCS(Å$^2$)
500
750
1000
m/z
20
0
−20
